# poRe GUIs for parallel and real-time processing of MinION sequence data

**DOI:** 10.1101/094979

**Authors:** Robert Stewart, Mick Watson

## Abstract

**Motivation:** Oxford Nanopore’s MinION device has matured rapidly and is now capable of producing over one million reads and several gigabases of sequence data per run. The nature of the MinION output requires new tools that are easy to use by scientists with a range of computational skills and which enable quick and simple QC and data extraction from MinION runs.

**Results:** We have developed two GUIs for the R package poRe that allow parallel and real-time processing of MinION datasets. Both GUIs are capable of extracting sequence- and meta- data from large MinION datasets via a friendly point-and-click interface using commodity hardware.

**Availability:** The GUIs are packaged within poRe which is available on SourceForge: https://source-forge.net/projects/rpore/files/. Documentation is available on GitHub: https://github.com/mw55309/poRe_docs

## 1 Introduction

Nanopore sequencing is the only sequencing technology that measures an actual single molecule of DNA, rather than incorporation events into a template strand ^1,2^. Early access to Oxford Nanopore’s MinION, a portable DNA sequencer approximately six inches in length, began in 2014. The MinION may be considered a mature platform, having been used to sequence bacterial genomes ^3,4^; resolve repeats in the human genome ^5^; study cDNA structure ^6,7^; detect base modifications^8–10^; detect antibiotic resistance ^11^; perform real-time enrichment (’read until’)^12^; and provide surveillance in a human disease outbreak^13^. The latest chemistry release, R9.4, has seen the first high-coverage human genome data released (https://github.com/nanopore-wgs-consortium/NA12878; https://github.com/nanoporetech/ONT-HG1), with several MinION flowcells from the two projects producing over 4 gigabases (Gb) of sequence data.

The MinION has been designed to enable mobile, real-time sequencing. As soon as a sequencing library is placed onto the device, the MinION begins sequencing. Each channel/nanopore reports asynchronously, creating a single file per channel per read. These are created in HDF5, a compressed binary hierarchical data format (https://www.hdfgroup.org/). Depending on the sequencer and chemistry version, these HDF5 files include raw or event-level signal data, recorded as a DNA molecule passed through the pore. There are a range of base-calling options, including cloud-based Metrichor, local MinKNOW base-calling and open-source alternatives ^14,15^, that will convert the signal data into DNA sequences.

The real-time nature of the platform (e.g. ^16^) is realised because both the signal-level and base-called HDF5 files are available within seconds/minutes (respectively) of a read being sequenced, which means that downstream pipelines can use the data almost immediately for analysis. This contrasts with other platforms which must wait until the sequencing run is finished before processing and analysing the data.

With 512 pores and a sequencing speed of 250 bases-per-second, each MinION flowcell has the capacity to produce several million reads in a 48-hour run. As each read presents as two files (one raw, one base-called) MinION runs represent huge challenges for researchers without sufficient computational skills. Tools exist, such as poRe ^17^, to assist with this, but many are command-line based, and there is a need for easy-to-use, GUI-based tools for MinION data QC and analysis.

## 2 Methods

We have designed and built two graphical-user-interfaces (GUIs) for MinON data processing, organization and extraction. Both are built as Shiny apps and released as part of the package poRe ^15^.

**Figure 1.**
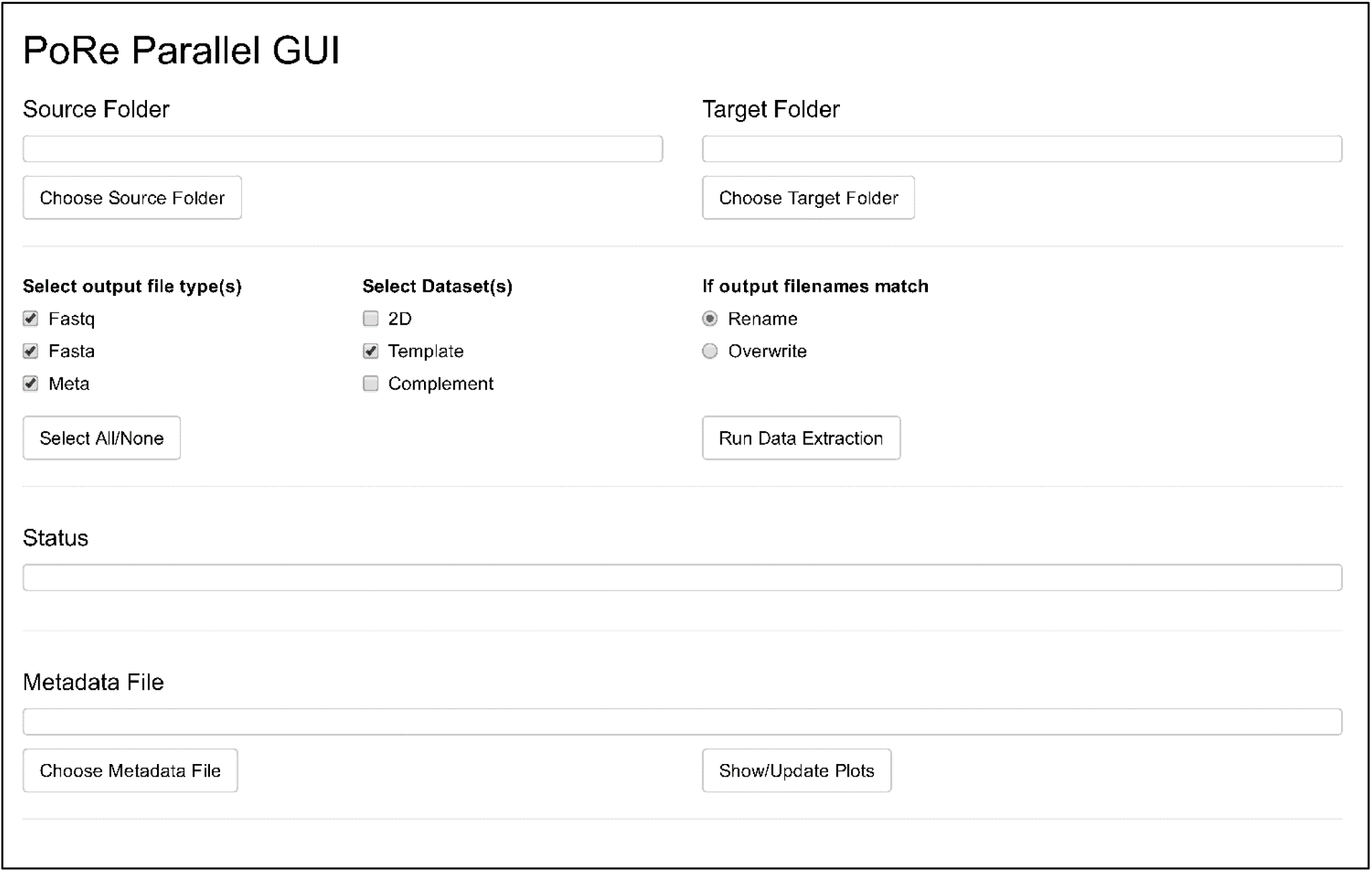
Screenshot of the pore parallel GUI, which as a Shiny App will open in the user’s browser.

The poRe real-time GUI is designed to extract data (FASTQ, FASTA and metadata) during a run, or during basecalling. A source and destination folder are required. The software then monitors the source folder for new FAST5 files; as FAST5 files arrive in the folder, they are processed, data are extracted and output to the destination folder. The poRe real-time GUI saves researchers a huge amount of time as data can be extracted while the MinION is running. The poRe real-time GUI has built in parallelization using R’s built in parallel package, and is accessed by running the command pore_rt().

The pore parallel GUI is designed to extract data from runs that have already finished. Again, the software expects a source and destination folder; in addition, the user can select which data to extract, and the number of cores to use. The software then extracts FASTQ, FASTA and metadata from all files in the source folder into files in the destination folder; using the number of cores specified by the user, via the parallel package. The poRe parallel GUI is accessed via the pore_parallel() command.

## 3 Results

The poRe parallel GUI was able to simultaneously extract FASTQ, FASTA and metadata from 209,819 FAST5 files downloaded from the “cliveome” project in just 37 minutes on our 16-core Linux server, at a rate of approx. 90 FAST5 files per second.

## Funding

This work was supported by The Biotechnology and Biological Sciences Research Council (BBSRC) including institute strategic support to The Roslin Institute (BB/M020037/1, BB/J004243/1, BB/J004235/1, BBS/E/D/20310000).

### Conflict of Interest

the authors have received free flowcells and reagents from Oxford Nanopore as part of the MAP. Mick Watson has attended Oxford Nanopore events and had his travel paid for by ONT.

